# Praxis-BGM: Clustering of Omics Data Using Semi-Supervised Transfer Learning for Gaussian Mixture Models via Natural-Gradient Variational Inference

**DOI:** 10.1101/2025.11.13.688299

**Authors:** Qiran Jia, Jesse A. Goodrich, David V. Conti

**Affiliations:** Division of Biostatistics and Health Data Science, Department of Population and Public Health Sciences, Keck School of Medicine, University of Southern California; Division of Environmental Health, Department of Population and Public Health Sciences, Keck School of Medicine, University of Southern California; Department of Biostatistics and Informatics, Colorado School of Public Health

**Keywords:** Bayesian Clustering, Statistical Transfer Learning, Variational Inference, Gaussian Mixture Model, Omics

## Abstract

**Motivation:** High-dimensional omics data are typically measured on limited sample sizes, which challenges model-based clustering methods such as Gaussian mixture models, often leading to instability and poor generalization under complex mixture structures. To address these limitations, we developed Praxis-BGM, a natural-gradient variational inference framework for Gaussian mixture models that enables semi-supervised transfer learning by incorporating an informative prior Gaussian mixture model derived from large-scale reference data with robust cluster structures. This prior can encode cluster-specific means, covariance structures, and structural connectivity patterns, and is updated using the target data with variational inference to improve clustering in small-sample settings.

**Results:** We derived natural-gradient updates for standard parameters and assess feature-level contributions to posterior clustering via Bayes Factors. Implemented in Python library JAX for accelerator-oriented computation, Praxis-BGM is computationally efficient and scalable. Across extensive simulations and two real-world applications—breast cancer bulk transcriptomics for subtype recovery and single-cell transcriptomics for cross-platform label transfer—Praxis-BGM improves posterior clustering performance, stability, and biological interpretability, even when priors are partially mismatched.

**Availability and Implementation:** Praxis-BGM is freely available at https://github.com/ContiLab-usc/Praxis-BGM, and an archival version is available on Zenodo at https://doi.org/10.5281/zenodo.19657680.

**Supplementary Information:** Supplementary materials are available with the manuscript submission.

## 1 Introduction

Omics data, such as transcriptomics, metabolomics, and proteomics, offer potential insight into human health and disease. Since omics data captures an enormous number of molecular parameters, the inherent patterns and complexities within these measurements can reveal distinct disease mechanisms, variations in disease risk, or heterogeneous survival probabilities [25, 33]. Unsupervised clustering methods, such as Gaussian Mixture Models (GMMs), infer underlying patterns and assign data points to latent clusters, playing a crucial role in omics analyses for data exploration, disease risk stratification, subtype discovery, cell-type identification, and other downstream inferences [12, 25]. However, complex correlations, high dimensionality, and limited sample sizes (HDLSS) in many cohort studies pose substantial challenges for statistical analysis [25, 33, 37, 43].

As a burgeoning concept, statistical transfer learning offers an effective strategy for addressing these challenges by leveraging a related source domain to improve modeling and inference in a target domain, especially in generalized linear model (GLM)-based settings [19, 27, 34]. Existing transfer-learning extensions of GMMs have mainly focused on unsupervised settings, in which all available datasets are jointly clustered. One example is a multi-task GMM framework that jointly learns multiple related GMMs by leveraging shared parameter structure across datasets and improving robustness to outlier datasets [36]. Distributed transfer GMMs, in contrast, learn node-specific clustering models while exchanging model information across decentralized sites to improve clustering without sharing raw data [38]. While useful, these approaches do not directly address the setting in which the source domain is well characterized by robust clusters, often with interpretable annotations, and a reference GMM can therefore be reliably fitted or constructed, such as when a large biobank or consortium dataset with cluster annotations is used to guide clustering in a smaller, more specialized clinical study. Bayesian inference provides a natural mechanism for transfer learning in this setting by encoding source-domain knowledge into informative priors [34, 39]. Accordingly, we propose Praxis-BGM, a semi-supervised Bayesian transfer-learning method for GMMs, in which the source-domain-derived GMM is used as an informative prior for inference in the target domain. Building on recent advances in natural gradient variational inference (NGVI) for mixtures of minimal exponential-family distributions [18, 21], Praxis-BGM effectively updates this prior model with target-domain data, allowing the source-informed prior model to regularize the posterior while still adapting to the observed target domain.

Figure 2 provides an overview of the Praxis-BGM workflow, illustrating prior construction from a source domain and NGVI-based update in the target domain. In this setting, the source domain is assumed to contain robust cluster structures or known curated cluster labels, and ideally to be annotated with meaningful biological interpretations (e.g., cancer subtypes). To the best of our knowledge, no existing Bayesian framework has been proposed for semi-supervised transfer learning in GMMs under labeled reference datasets. This provides a principled transfer mechanism that complements existing GLM-based transfer learning strategies [19]. We also showed that priors can be constructed not only from a source dataset in the same feature space with different samples, but also from an auxiliary omic layer measured on the same samples, thereby enabling multi-omics in-serial integration, as explored in previous studies but within different modeling frameworks [14, 43].

**Figure 2:**
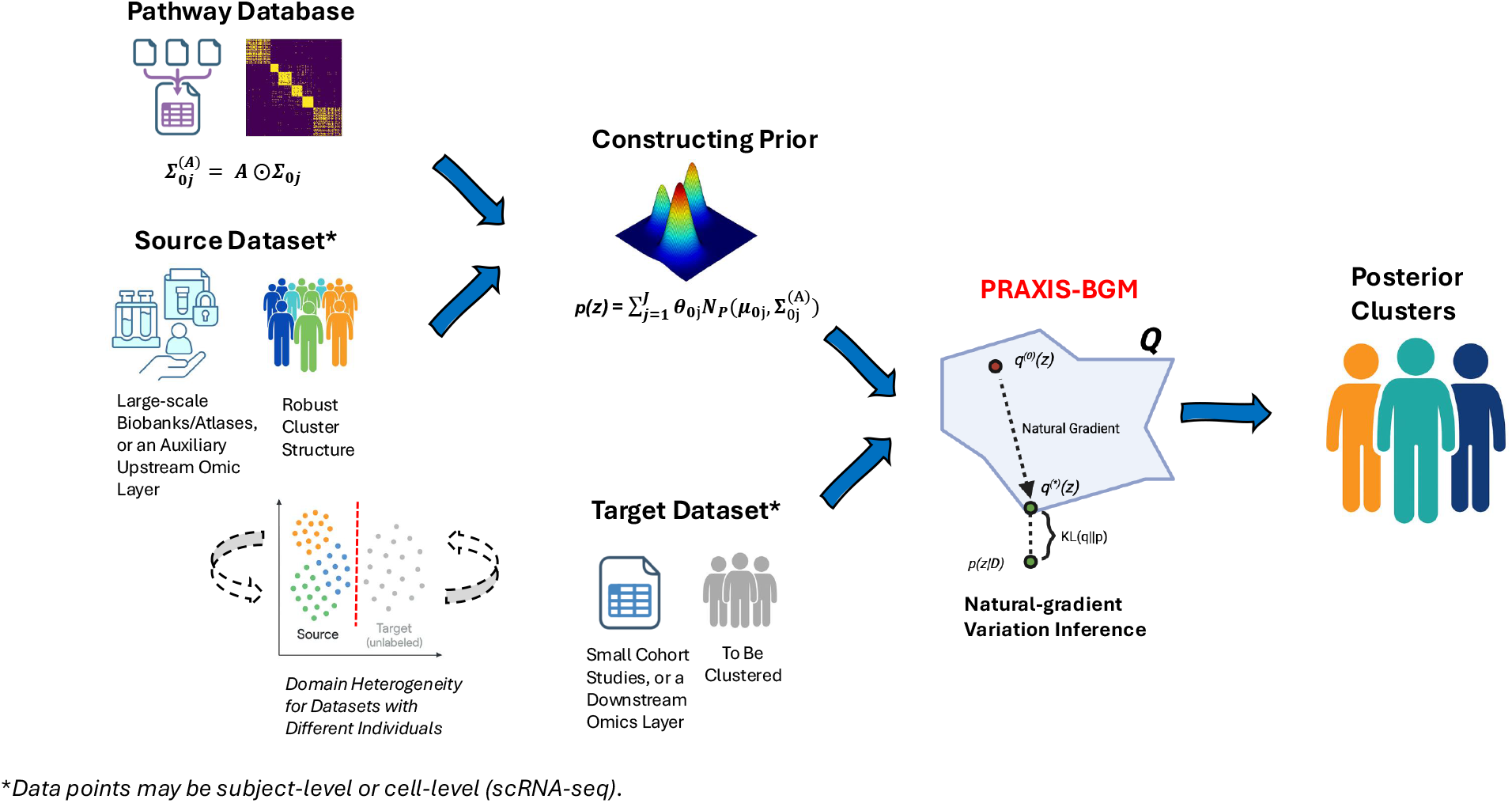
Overview of the Praxis-BGM framework for prior construction and natural-gradient variational inference-based Bayesian transfer learning for Gaussian Mixture Models, where the source of prior information may originate either from a different dataset measured on different samples (e.g., large-scale biobanks or atlases) or from an informative upstream omic layer measured on the same samples (e.g., from RNA-seq to miRNA).

We applied Praxis-BGM to breast cancer subtyping across transcriptomic datasets to transfer subtype structure from a reference cohort to a target cohort, and subsequently performed in-serial integrative clustering using Praxis-BGM from RNA-seq to miRNA within the same subjects to refine molecular subtypes. We also applied the method to the task of label transfer across single-cell RNA-seq technologies (i.e., sequencing platforms), such as inDrop (droplet-based) and CEL-seq, using pancreatic single-cell RNA-seq data, where transfer and inference were performed in the latent embedding space. In both applications, ground-truth labels were either fully or partially available for the query data, allowing for a direct validation of Praxis-BGM’s superior performance in transferring prior knowledge and guiding clustering.

The remainder of the paper is structured as follows. Section 2 details the proposed method and optimization with the NGVI framework. Section 3 introduces the real-world data and outlines the experimental settings for applications. Section 4 reports the results, demonstrating the strong and consistent advantages of Praxis-BGM. We also present extensive simulation results evaluating the effectiveness of Praxis-BGM in transferring cluster structure via individual components of informative priors, as well as jointly, across diverse signal-to-noise regimes and source–target discrepancies in Supplementary Data S.1 and S.2. Overall, these simulation results demonstrate that Praxis-BGM achieves efficient, robust and consistently superior performance relative to competing methods, even under challenging and partially misspecified conditions. Finally, Section 5 concludes with a discussion of our findings and future research directions.

## 2 Methods

Let *z* ∈ ℝ^*P*^ be a random variable following a GMM, i.e., a mixture of multivariate Gaussian distributions, with *J* components. Its marginal density is

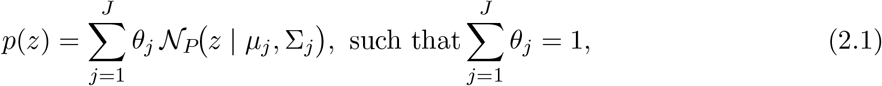

where *θ*_*j*_ is the mixing weight for component *j, µ*_*j*_ ∈ ℝ^*P*^ is the corresponding mean vector, and 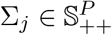 is the corresponding symmetric positive definite (SPD) covariance matrix.

### 2.1 Priors from Source Data in the Same Feature Space as the Target Data

Suppose a source dataset *D*_0_, measured in the same *P*-dimensional feature space as the target dataset *D*, is available from a related population. We leverage *D*_0_ to construct informative priors for *z*. This yields a Bayesian transfer-learning formulation in which the source data inform the prior distribution *p*(*z* | *D*_0_), and the target data *D* updates this prior through Bayes’ rule [34]. Specifically,

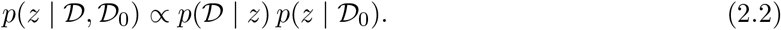

The normalizing constant *p*(*D* | *D*_0_) = *p*(*D* | *z*) *p*(*z* | *D*_0_) *dz* is generally intractable. Therefore, in a two-stage procedure, we first estimate the source-informed prior distribution *p*(*z* | *D*_0_) and then employ variational inference to approximate the posterior *p*(*z* | *D*, *D*_0_) by maximizing a tractable lower bound on the log marginal likelihood, namely the evidence lower bound (ELBO) [4].

Specifically, we parameterize the prior *p*(*z* | *D*_0_) as a GMM with parameters 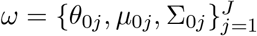, representing the distribution learned from *D*_0_. When *D*_0_ includes cluster labels **X**_0_, we can directly use cluster-specific summary statistics or discriminant analysis to derive *ω*. In the absence of labels, a standard GMM can be used to estimate the prior mixture distribution, with the optimal number of clusters selected using standard criteria such as the Bayesian Information Criterion (BIC) [12, 37]. It is often useful to inflate the variance of the source-informed prior, as the distribution of *z* may differ substantially across domains, to improve robustness [34]. Specifically, for each component *j*, we replace Σ_0*j*_ by *c* Σ_0*j*_ with *c >* 1. The prior cluster-specific covariance matrices Σ_0*j*_ can optionally be adjusted by additional prior knowledge to encourage sparsity as

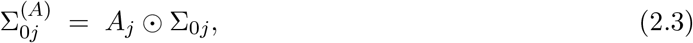

where the sparsity-imposing matrix *A_j_* ∈ {0, 1} ^*P ×P*^ is symmetric with unit diagonal (*A*_*j,p p*_ = 1, ∀ *p*), and is derived from grouping information, such as that available from existing pathway databases to encode element-wise structural modulation [35]. The matrix *A*_*j*_ shapes the prior covariance matrix via a Hadamard mask as shown in Equation (2.3).

### 2.2 Priors from an Auxiliary Omic Layer Measured on the Same Samples as the Target Data

We also allow for constructing informative priors from an auxiliary or multi-omic layer collected on the same samples as *D*. Let *D*^(*O*)^ denote an alternative omic layer (e.g., DNA methylation) measured on the same subjects who provide the target data *D* (e.g., transcriptomics). When cluster or class labels **X**^(*O*)^ are available or can be reliably inferred from *D*^(*O*)^, we construct a prior distribution *p*(*z* | *D*^(*O*)^) parameterized by 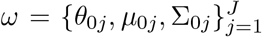, where the parameters are obtained from discriminant analysis or empirical class-conditional summary statistics in the target feature space [12]. This enables using a biologically related but distinct omic layer, measured on the same subjects, to derive the prior distribution for the latent variable *z* representing upstream cluster-structure signals, in line with in-serial integration strategies for multi-omics data [14, 43].

### 2.3 Variational Inference with Informative Priors

Given informative priors derived from either the source data *D*_0_ or the auxiliary omic layer *D*^(*O*)^, denoted collectively by *p*_*ω*_(*z*) for simplicity, we approximate the posterior with a surrogate finite mixture of multivariate Gaussian distributions *q*_*ϕ*_(*z*) with 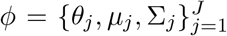 using a NGVI framework [18, 21]. Given priors *p*_*ω*_(*z*) and target data *D*, we seek a variational distribution that maximizes the ELBO,

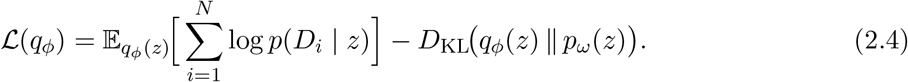

Maximizing ℒ(*q*_*ϕ*_) encourages *q*_*ϕ*_(*z*) to explain the target mixture data *D* well, while remaining regularized toward the prior *p*_*ω*_(*z*). In general, NGVI is a first-order optimization method for maximizing ℒ(*q*_*ϕ*_) of minimal exponential-family (EF) distributions that replaces the standard Euclidean gradient with the natural gradient, thereby accounting for the information geometry of the natural parameter space [18]. Natural gradient yields parameterization invariance and often significantly improves convergence efficiency and stability [1, 18]. Specifically, NGVI updates the natural parameter ***λ***_*z*_ by scaling the standard Euclidean gradient with the inverse Fisher information matrix (FIM): 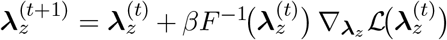, where *β* is the step size. Direct computation and inversion of the FIM is computationally expensive in high-dimensional settings. For minimal EF variational distributions, one can avoid this explicit evaluation by working in the expectation-parameter space ***m***_*z*_, using the duality between natural and expectation parameters [18]. Under this duality, the natural gradient for ***λ***_*z*_ can thus be expressed as the ordinary gradient with respect to 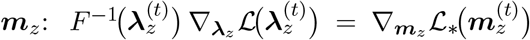, where ℒ_*_ denotes the ELBO expressed in the expectation-parameter space. 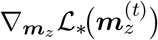 can be obtained in terms of gradients with respect to the standard parameters (e.g., ***µ*** and **Σ** for Guassian) by using the chain rule, which in turn yields the corresponding natural-gradient update for standard parameters via the variational online Newton (VON) algorithm [21]. NGVI has been extended beyond minimal EF to mixtures of minimal EF distributions, including finite mixtures of multivariate Gaussians (i.e., GMMs), under a minimal conditional EF representation [21, 24].

We apply NGVI for GMMs with informative priors, *p*_*ω*_(*z*), to facilitate Bayesian transfer learning. We define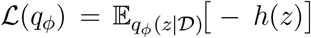, and 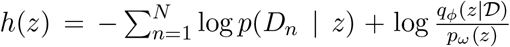. The VON algorithm is used to efficiently evaluate *h*(*z*) by drawing a stochastic sample under *q*_*ϕ*_(*z*) and then use it to compute natural gradients for standard parameters *ϕ* of GMMs [18, 21]. Specifically, at iteration *t*, we first draw a component *v* and then draw a sample conditioned on the component,

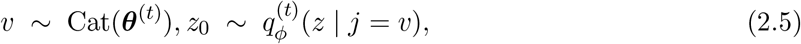

which serves as the anchor point for natural-gradient updates. We then define

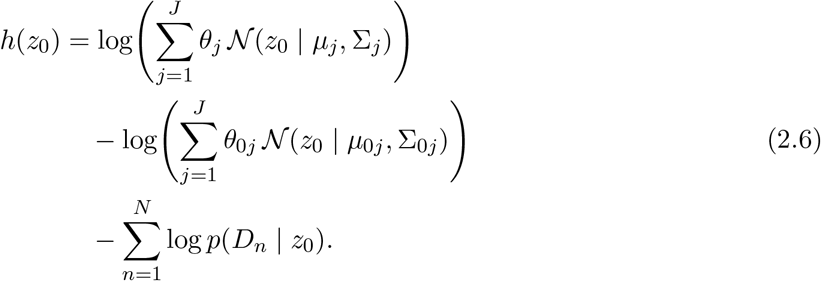

The stochastic gradient and Hessian of *h*(*z*_0_) with respect to *z*_0_, denoted by ∇_*z*_*h*(*z*_0_) and 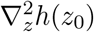, respectively, are key quantities in the derived NGVI updates for standard parameters *ϕ*, obtained using Bonnet’s and Price’s theorems as shown in [21]:

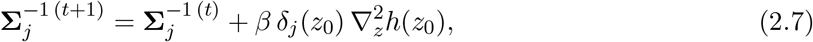

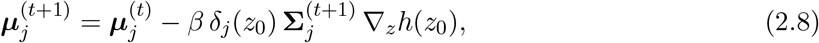

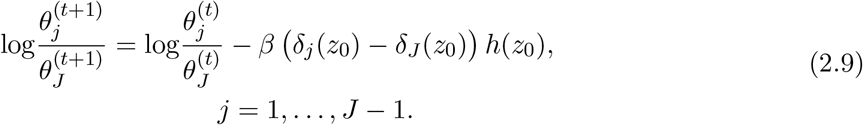

where

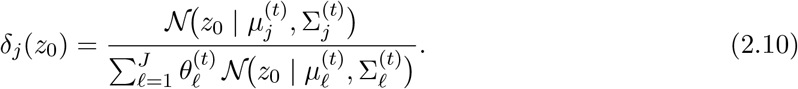

We summarize Praxis-BGM (prior-augmented NGVI for GMMs) in Algorithm 1. We initialize the variational parameters *ϕ* using the prior parameters *ω* to facilitate alignment between the target-domain cluster labels and the prior structure, thereby enhancing interpretability. Praxis-BGM is implemented in Python using JAX, enabling accelerator-friendly computation on GPUs and TPUs, as well as just-in-time compilation for scalable and numerically stable NGVI [5].

#### Algorithm 1.

Praxis-BGM: Prior-augmented Natural Gradient Variational Inference for Gaussian Mixture Models

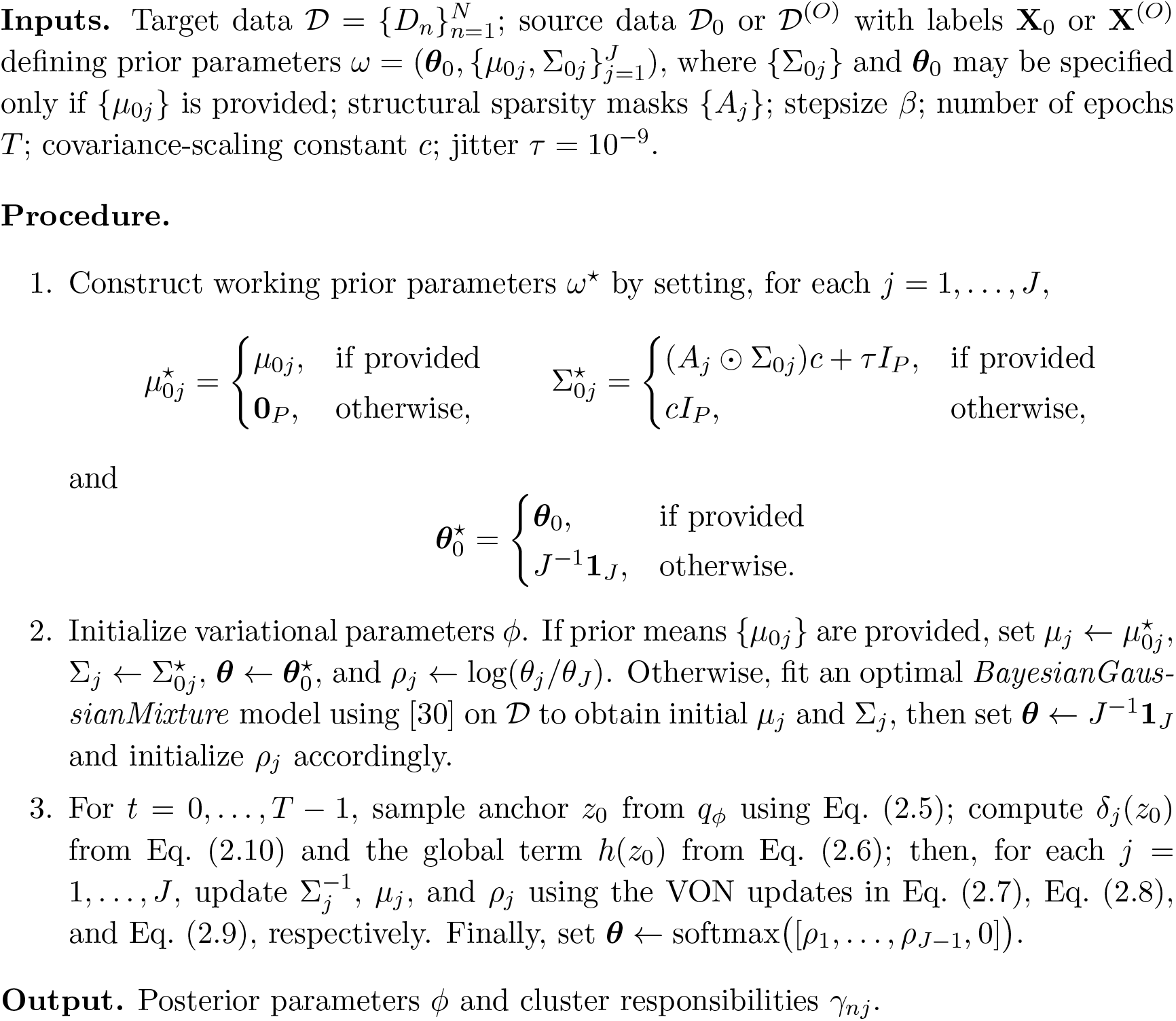

## 3 Evaluation with Real-world Data

We demonstrated the effectiveness of Praxis-BGM in transferring prior cluster structural information to improve clustering performance and enhance clinical relevance compared with standard clustering and classification methods across two real-world data applications.

### 3.1 Breast Cancer Bulk Transcriptomics

The Cancer Genome Atlas Breast Invasive Carcinoma (TCGA-BRCA) and the Molecular Taxonomy of Breast Cancer International Consortium (METABRIC) are two breast cancer genomic cohorts [9, 26], each profiling the transcriptomic expression of ∼ 20,000 genes (bulk RNA-seq data) and accompanied by clinical and survival information. TCGA-BRCA includes PAM50 subtype labels for 500 subjects, whereas METABRIC provides hormone-receptor status, a well-established surrogate strongly correlated with intrinsic subtypes [28, 31]. For METABRIC, we constructed surrogate subtype labels by mapping clinical receptor status to their closest intrinsic subtype counterparts. Specifically, ER+/HER2− cases were categorized as Luminal A-like, ER+/HER2+ as Luminal B-like, ER−/PR−/HER2+ as HER2-enriched-like, and ER−/PR−/HER2− as Triple-negative, used here as a surrogate for Basal-like subtype given their strong overlap [31]. These labels are interpreted as approximate clinical proxies rather than definitive molecular subtype assignments, and we used them to derive related but not fully accurate informative priors for transfer-learning-based clustering. For TCGA-BRCA, we utilized the RNA-based PAM50 labels available as ground-truth, noting that 560 subjects (52.83%) in the expanded cohort lack these formal molecular annotations.

The RNA-seq data of METABRIC was used as the labeled source dataset to derive priors for Praxis-BGM, with the RNA-seq data of TCGA-BRCA serving as the unlabeled target dataset. We hypothesized that incorporating informative priors in METABRIC would yield clusters in TCGA-BRCA that better align with PAM50 subtypes and exhibit stronger prognostic relevance than clustering or supervised prediction across datasets alone. After filtering incomplete records, 1,980 METABRIC and 1,060 TCGA-BRCA samples remained. Using log_2_ transformed gene expressions shared across datasets, we selected the top 1,000 most variable genes based on METABRIC based on an elbow-based selection method on the ranked gene variance plot to remove low-information genes. We then applied standardization and subset both datasets accordingly. For each METABRIC subtype, we computed empirical means and covariance matrices to form component-specific priors for Praxis-BGM.

For performance evaluation, Praxis-BGM clusters were compared with the unsupervised Bayesian Gaussian Mixture Models (BGM) [37], a VI-based GMM method with uninformative priors (fit with *J* = 4), and supervised classifiers trained on the source data and applied to the target data (multinomial logistic regression, Linear Discriminant Analysis (LDA) [12], and Extreme Gradient Boosting (XGBoost) [7] tuned via cross-validation). Estimated cluster labels in TCGA-BRCA were evaluated by Adjusted Rand Index (ARI) and accuracy with PAM50 labels, and were further evaluated as predictors in separate age-adjusted Cox proportional hazards models for five-year survival. Bayes factors (BFs) for feature importance in driving breast cancer subtype differentiation (method detailed in Supplementary Data S.3.1) were computed to identify subtype-driven genes, followed by functional enrichment analysis using Gene Ontology (GO) Biological Process [2, 8] and Human Molecular Signatures Database (MSigDB) Hallmark gene sets [20] via `GSEApy`[11].

We next propagated the estimated RNA-seq Praxis-BGM clusters as priors for clustering of the microRNA (miRNA) layer in the same TCGA-BRCA cohort, enabling a second Bayesian update of the GMM while achieving in-serial multi-omics integration [14, 43]. Biologically, miRNA captures regulatory signals not fully reflected by RNA-seq expression alone and may provide complementary information for breast cancer subtyping [3]. This in-serial analysis follows Section 2.2 and the analysis method is detailed in Supplementary Data S.3.4.

### 3.2 Cross-platform Label Transfer in Pancreatic scRNA-seq

When analyzing single-cell RNA sequencing (scRNA-seq) data, a key initial task is to annotate cell types for clusters of cells. The increasing availability of annotated reference datasets has led to numerous label-transfer methods [29]. Among them, scANVI [40], a semi-supervised extension of scVI [22], is a state-of-the-art approach that jointly models labeled reference and unlabeled query data. The `Scanpy`framework [15] also provides K-Nearest Neighbors(KNN)-based label-transfer functionality by integrating PC embeddings across datasets, called `ingest`, and heterogeneity-correction methods, such as ComBat-seq [42] and Scanorama [16], can be paired with KNN classifiers for label transfer.

Label transfer naturally aligns with the goal of Praxis-BGM, which performs clustering guided by cell-type-specific priors. We used the human pancreas scRNA-seq atlas assembled by [23], a benchmark dataset of 16,382 cells annotated into 14 cell types across four platforms (inDrop, CEL-Seq, Smart-seq, Fluidigm C1), chosen for its heterogeneous batch structure. After quality control to remove low-quality cells (zero-count entries), we applied log_2_ transformation, selected the top 2,000 highly variable genes (HVGs) using `Scanpy`, standardized them, performed PCA, and retained the top 100 principal components. Cells from inDrop (52.3%) served as the reference dataset, while cells from the remaining platforms (47.7%) formed the query dataset. Ground-truth annotations were available for all cells and used only for evaluation on the query subset. Figure 4(a) shows the cell-type distributions across reference and query sets.

For Praxis-BGM, we computed cell-type-specific mean priors in the PC space corrected by ComBat-seq from the reference data and applied them to guide clustering on the query data. We benchmarked Praxis-BGM against scANVI, `Scanpy`‘s `ingest`function, and integration-based methods (ComBat-seq and Scanorama) followed by KNN classification. Notably, Praxis-BGM and scANVI use the reference information differently: Praxis-BGM incorporates priors directly into cluster inference on the query set, whereas scANVI learns a shared latent embedding for both datasets before label prediction. For computational time comparisons, we measured only the cost of heterogeneity correction and inference, excluding embedding computation.

## 4 Results

### 4.1 Praxis-BGM Enhances Breast Cancer Subtype Recovery and Clinical Relevance in TCGA-BRCA

Our results show that, by leveraging informative priors derived from METABRIC, Praxis-BGM improved both subtype concordance and clinical interpretability relative to unsupervised clustering and supervised classification approaches applied to TCGA-BRCA RNA-seq data.

Table 1 presents the ARI and accuracy results for concordance with ground-truth PAM50 subtype labels for Praxis-BGM and other benchmark methods on TCGA-BRCA RNA-seq data. BGM clusters captured partial concordance with the PAM50 molecular subtypes (ARI = 0.392), indicating that even without subtype-specific priors, the intrinsic transcriptomic structure of TCGA-BRCA samples reflects moderate underlying subtype-related patterns. For classification benchmark methods, multinomial regression and LDA produced similar and better subtype concordance (ARI = 0.381 and 0.379, accuracy = 0.654 and 0.650 respectively). XGBoost yielded the lowest ARI (0.351) and accuracy (0.638) among the reported supervised classification methods, suggesting potential overfitting despite the use of 10-fold cross-validation for hyperparameter tuning. Among all methods, Praxis-BGM achieved the strongest overall performance (ARI = 0.430; accuracy = 0.714). We then visualized the 5-year survival across Praxis-BGM clusters using Kaplan Meier curves in Panel (a) Figure 3. The Kaplan–Meier curves show clear survival heterogeneity across the inferred Praxis-BGM clusters. The posterior Praxis-BGM cluster corresponding to Luminal A has the best prognosis, whereas the cluster corresponding to Luminal B (HER2+) has the poorest survival with the steepest decline. Overall, Praxis-BGM clusters yield the strongest survival stratification under the age-adjusted Cox proportional hazards model for 5-year survival (LRT statistic = 57.79, p-value = 8.460 × 10^−12^). The Cox model using BGM clusters is likewise significant (LRT statistic = 45.121, p-value = 3.752 × 10^−9^) but less pronounced, whereas all three classification methods show weaker significance in their corresponding Cox models. Taken together, these findings show that Praxis-BGM clusters not only recover established subtype structure, but also provide superior prognostic stratification beyond benchmark unsupervised and supervised methods through Bayesian transfer learning. As for runtime, although BGM achieved the fastest CPU runtime, Praxis-BGM remained computationally efficient and benefited from TPU acceleration.

**Table 1:** Benchmarking clustering and classification methods on TCGA-BRCA RNA-seq data by assessing the ARI and accuracy with ground-truth PAM50 subtype labels, restricted to the 500 subset of samples with available subtype annotations, and by evaluating prognostic value using the likelihood-ratio test (LRT) statistic and corresponding p-value from age-adjusted Cox proportional hazards models for 5-year survival, and reporting runtime with the corresponding hardware.

| Method | ARI | Accuracy | LRT Stat (p-value) | Time (s) / Hardware |
| --- | --- | --- | --- | --- |
| Praxis-BGM | **0.430** | **0.714** | **57.787** ($8.460 \times 10^{-12}$) | 6.04 /V6e TPU |
| BGM | 0.392 | - | 45.121 ($3.752 \times 10^{-9}$) | **1.77**/CPU |
| Multinomial Regression | 0.381 | 0.654 | 31.711 ($2.191 \times 10^{-6}$) | 2.24/CPU |
| LDA | 0.379 | 0.650 | 33.661 ($8.744 \times 10^{-7}$) | 11.08/CPU |
| XGBoost | 0.351 | 0.638 | 30.556 ($3.771 \times 10^{-6}$) | 11.82/CPU |

**Figure 3:**
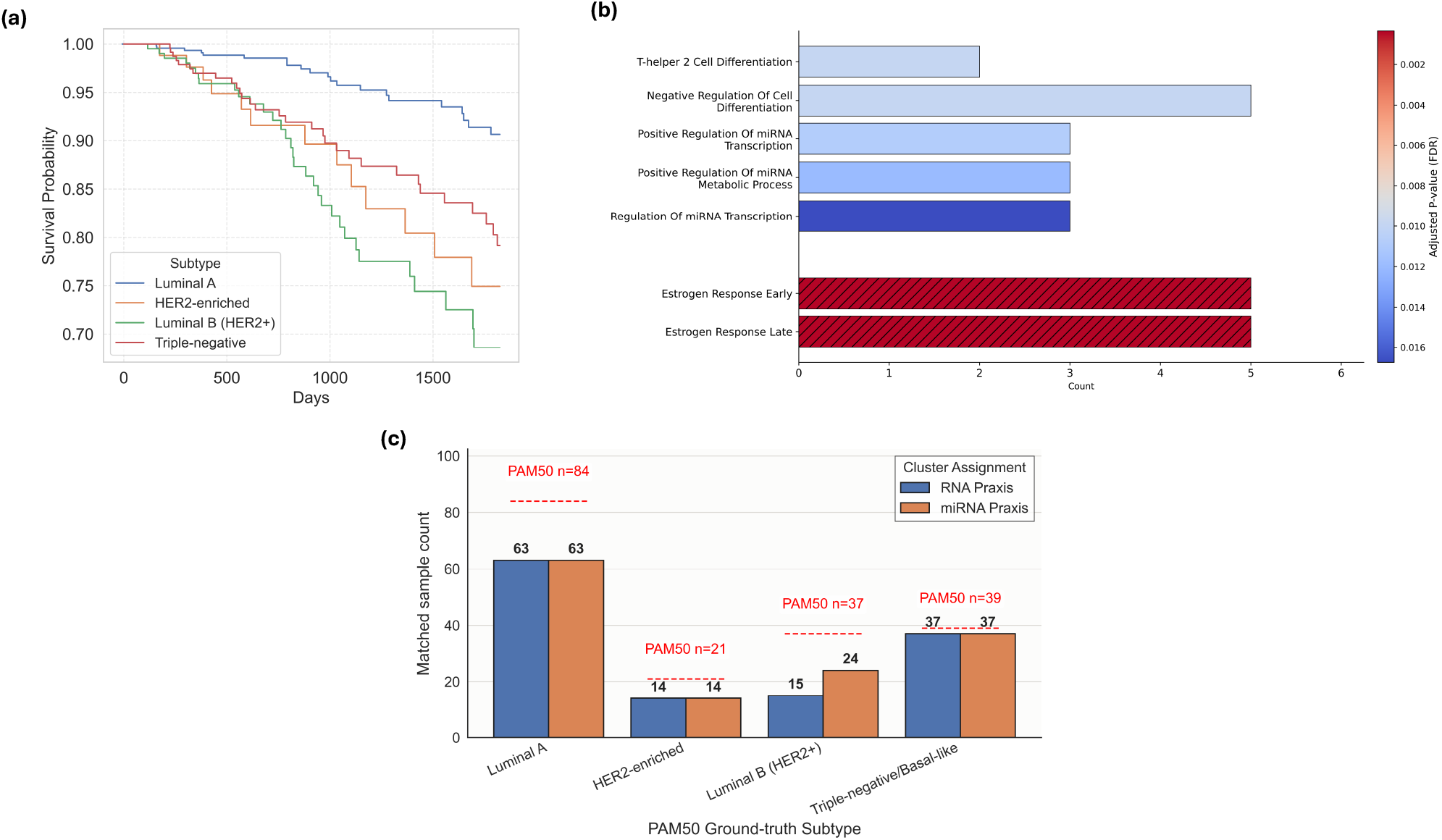
(a) Kaplan–Meier curve comparing survival across Praxis-BGM clusters annotated by subtypes; (b) Pathway enrichment analysis of the non-indeterminate genes (BF *>* 10^0.5^). The solid bars represent pathways from the Gene Ontology (GO) Biological Process database, whereas the hatched bars represent Human Molecular Signatures Database (MSigDB) Hallmark gene sets; (c) Subtype-specific concordance of RNA- and miRNA-based Praxis-BGM clusters with PAM50 ground-truth labels.

To identify genes driving breast cancer subtype differentiation, we computed BFs for all genes and identified 39 genes showing at least substantial evidence based on Jeffreys’ scale [17]. To better interpret the identified clustering-drive genes, Panel (b) of Figure 3 shows the results of the pathway over-representation analysis for genes with at least substantial evidence (BF *>* 10^0.5^). After FDR adjustment, seven pathways were significant. Interestingly, the two estrogen response pathways found in MSigDB are well-established signatures in the progression of breast cancer [41]. Among GO Biological Process, the three miRNA-related significant pathways are also biologically relevant to breast cancer, as aberrant miRNA expression has been implicated in tumor initiation, progression, and therapeutic resistance [3]. These results suggest that integrating informative priors from METABRIC through Praxis-BGM not only yields more clinically relevant clusters, but also improves biological interpretability and downstream analysis, further motivating the in-serial integration of the miRNA layer in the target data.

For the in-serial analysis with miRNA data in TCGA-BRCA, leveraging the posterior RNA-seq Praxis-BGM clusters as priors, the posterior miRNA Praxis-BGM clusters were largely consistent with the RNA-based Praxis-BGM clusters while providing refined stratification that improved alignment with PAM50 labels among subjects with available annotations and miRNA data. Specifically, clustering agreement measured by ARI increased from 0.417 to 0.478. To explore this increase, in Panel (c) of Figure 3 we presents the subtype-specific concordance of the RNA-and in-serial miRNA-based Praxis-BGM clusters with PAM50 ground-truth labels. Notably, the Praxis-BGM clusters updated with the miRNA layer correctly assigned 9 additional subjects to the Luminal B subtype compared with the RNA-based Praxis clusters. These refined posterior miRNA Praxis-BGM clusters incorporate priors from both the external source dataset METABRIC and the RNA-seq layer of TCGA-BRCA, providing an illustrative example of multi-omics integration through sequential Bayesian updating.

### 4.2 Praxis-BGM Outperforms Existing Label Transfer Methods on Cross-platform scRNA-seq

Benchmarking on cross-platform scRNA-seq data revealed that Praxis-BGM provides superior label transfer accuracy and robustness compared with widely used alternatives.

Table 2 presents the results of benchmark label transfer methods on the query subset. Praxis-BGM and ComBat-seq + KNN substantially outperformed other methods in terms of ARI and accuracy, while also being computationally faster. Notably, Praxis-BGM achieved the highest clustering accuracy and ARI (accuracy = 0.975 and ARI = 0.957), better than those of ComBat-seq + KNN and scANVI. Because ComBat-seq + KNN uses the same corrected PC embeddings as Praxis-BGM, the 2.2% improvement in accuracy and 0.024 improvement in ARI further highlight the performance advantage of Praxis-BGM over the deterministic classification method of KNN. `Scanpy`-`ingest`and Scanorama + KNN yielded inferior ARI and accuracy, and required longer computation times than Praxis-BGM did. Among all methods, scANVI was the most computationally intensive as it is neural-network-based.

**Table 2:** Benchmarking label transfer methods on Pancreas single-cell transcriptomic data against expert annotations, and reporting runtime with the corresponding hardware platform.

| Method | ARI | Accuracy | Time (s) | Hardware | Embedding |
| --- | --- | --- | --- | --- | --- |
| Praxis-BGM | **0.957** | **0.975** | **2.94** | T4 GPU | PCs |
| ComBat-seq + KNN | 0.933 | 0.953 | 3.4 | CPU | PCs |
| scANVI | 0.912 | 0.921 | 33.0 | T4 GPU | scVI |
| Scanorama + KNN | 0.913 | 0.916 | 15.5 | CPU | PCs |
| Scanpy-ingest | 0.896 | 0.915 | 10.3 | CPU | PCs |

Praxis-BGM and scANVI relied on different embeddings to perform the label transfer (PCs for Praxis-BGM and scVI representations for scANVI). The improved performance of Praxis-BGM compared with other KNN-based methods that also use PC embeddings suggests a potential advantage of Praxis-BGM in better adapting to settings where the reference and query domains are not fully homogeneous leveraging variational inference. Figure 4(b) provides the confusion heatmap comparing predicted cell types to ground-truth annotations of the cell types in the query data. No-tably, the annotated reference subset contained 7 T cells while the query subset did not (Fig. 4(a)). ComBat-seq + KNN and scANVI exhibited pronounced misassignment of Acinar cells to Ductal, while Praxis-BGM did not. All methods showed some degree of confusion between Quiescent and Activated Stellate cells, likely due to their similarity, and all performed somewhat less accurately on the rarer cell types of Mast and Epsilon. Praxis-BGM and scANVI accurately assigned the rare cell type of Schwann, while ComBat-seq + KNN did not. In addition, we conducted an additional sensitivity analysis investigating whether KNN-based benchmark label-transfer methods would benefit from using a smaller and more targeted set of source-derived differentially expressed genes (DEGs) in Supplementary Data S.4.1, rather than the original workflow based on 2,000 HVGs. Overall, more aggressive DEG restriction modestly improved KNN-based benchmark methods, while Praxis-BGM remained the best-performing approach across all methods. We also evaluated a scenario in which the reference dataset contained fewer rare cell types than the query dataset (Supplementary Data S.4.2). Under this mismatch, Praxis-BGM still outperformed the benchmark methods.

**Figure 4:**
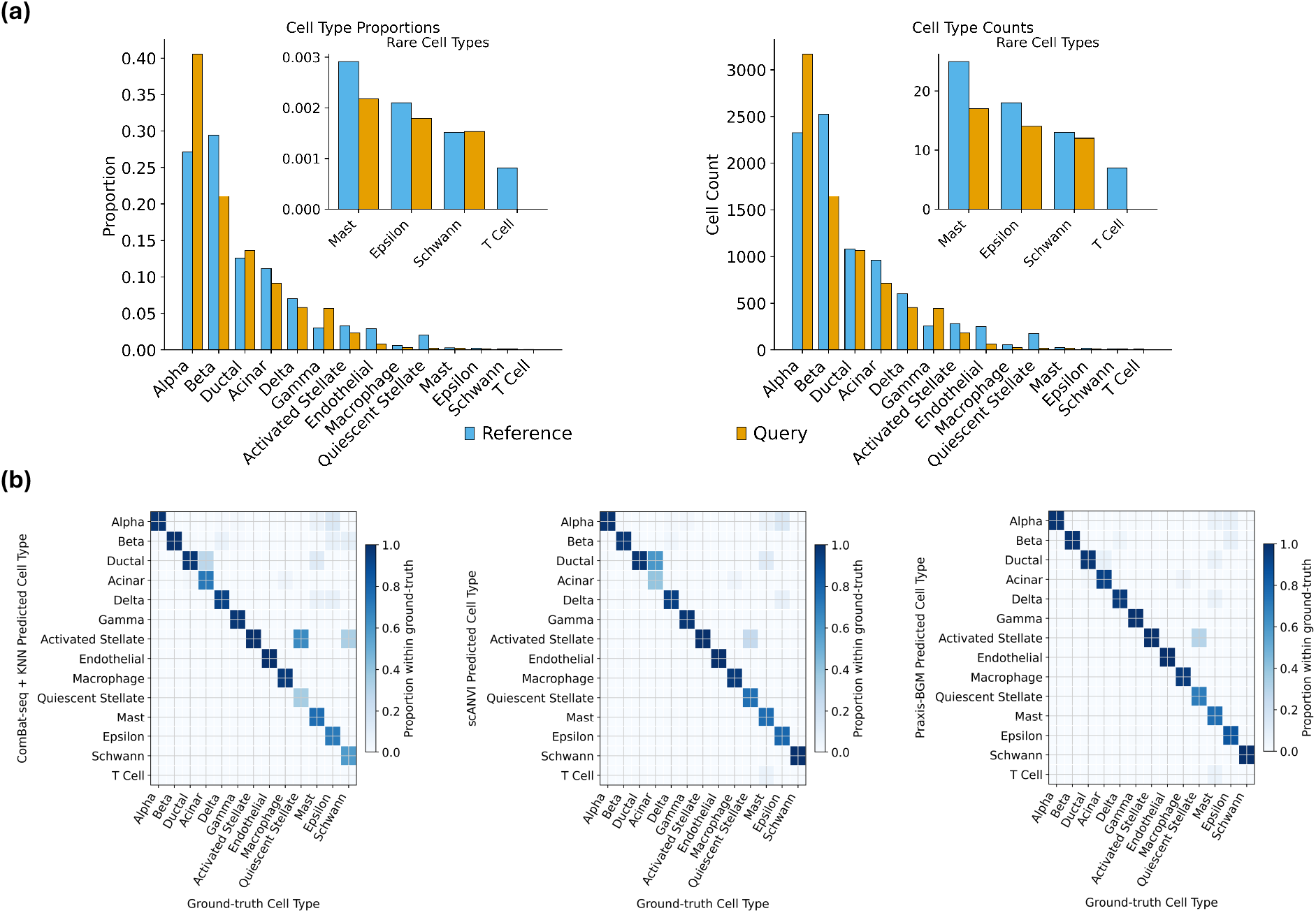
(a) Ground-truth cell type proportions (left) and count (right) distributions for the reference and query datasets. (b) Confusion heatmap comparing predicted cell types to ground-truth annotations in the query data for ComBat-seq + KNN, scANVI, and Praxis-BGM, with color intensity representing the proportion within each ground-truth cell type.

This application demonstrates that Praxis-BGM can leverage cell-type-informative priors to achieve accurate and computationally efficient label transfer between heterogeneous single-cell datasets, making it a practical and scalable alternative to existing methods. Looking ahead, as emerging technologies (e.g., in situ spatial transcriptomics) enable more comprehensive characterization of rare cell types, increasingly comprehensive reference datasets can be used for Praxis-BGM. Capable of addressing platform heterogeneity, Praxis-BGM is also especially useful for transferring well-annotated scRNA-seq labels to spatial RNA-seq data, where direct cell-type identification remains challenging.

## 5 Discussion

In this paper, we introduced Praxis-BGM, a semi-supervised GMM clustering method based on NGVI that facilitates transfer learning. The source data are assumed to have curated cluster labels, which, if not already available, can be obtained by fitting an unsupervised GMM with an appropriate model selection procedure and subsequently annotating the labels through biological interpretation. Relying on the source data, Praxis-BGM provides a highly flexible framework for incorporating prior information through a prior GMM estimated and updating the prior model with the target data through variational inference. One or any combination of cluster-specific parameters in the prior GMM can be specified, including the mean vector (prior belief in differences in expression levels across clusters), covariance matrix (prior belief in differences in variability and correlation patterns across clusters), and weights (prior belief in differences in cluster prevalence). In addition, overall or cluster-specific sparsity-inducing matrices informed from pathway and ontology databases can be incorporated to encode feature connectivity further. These prior components define a prior GMM and regularize the posterior estimation of the GMM on the target data. We demonstrated the superior performance of Praxis-BGM over competing deterministic classification-based methods in both simulation studies and real-world applications, as well as the advantage of incorporating prior knowledge over purely unsupervised clustering. In the context of breast cancer, Praxis-BGM successfully transferred subtype-specific component structures from METABRIC to TCGA-BRCA, yielding significant improvements in biological relevance and clinical utility while also enabling the discovery of cluster-associated genes. We further showed that Praxis-BGM can flexibly integrate multi-omic layers for in-serial analysis by using cluster-specific priors derived from a preceding omic layer to inform the analysis of the next layer. We also demonstrated that the application of Praxis-BGM extends naturally to label transfer in single-cell transcriptomics to effectively leverage annotated reference datasets to cluster and annotate query cells.

When multiple omics layers are available, the choice of which layer to prioritize for transfer learning, and whether a downstream in-serial update with additional omics layers is beneficial, depends mainly on two considerations: (i) biological relevance to the scientific question and study objective, and (ii) data availability and completeness. For example, in the breast cancer bulk transcriptomics application, our primary objective and main validation metric were to recover PAM50 labels in the target data. Accordingly, the transcriptomic layer is naturally the most informative layer for this objective, whereas the miRNA layer may offer complementary refinement for explorative analysis. By contrast, other omics layers, such as copy number variation (CNV), may not be prioritized for this specific goal. In practice, it is also often reasonable to begin with the layer measured in the largest and most complete set of subjects. More broadly, Praxis-BGM is a highly flexible approach, as priors may be derived from an external source dataset in the same feature space, from another omics layer measured in the same cohort, or from a sequential combination of both, as illustrated in the breast cancer bulk transcriptomics application. The most appropriate choice of modeling strategy depends on the research objective, the underlying biology, and the structure and availability of the source and target data. Moreover, when the source dataset contains high-quality integrative cluster labels estimated from multi-omics data, Praxis-BGM can still leverage that information to perform transfer learning using a single omics layer shared between the source and target datasets; the target dataset does not need to contain all omics layers available in the source data. An important direction for future work is the development of a parallel multi-omics transfer-learning framework that jointly integrates multiple omics layers while allowing layer-specific priors to be derived from potentially distinct source datasets.

One caveat about statistical transfer learning is the risk of negative transfer, which could occur when the knowledge transferred from the source data reduces the performance of modeling and inference in the target data [27]. Inappropriate priors from unsuitable source data for Praxis-BGM can also potentially lead to negatively biased posteriors. Our Variational Bayes framework seeks to mitigate this potential negative impact by reconciling discrepancies between priors and data through variational inference. Simulation studies demonstrate that Praxis-BGM maintains strong performance even under severely mis-specified priors. Future work could focus on enhancing robustness by learning the reliability of priors derived from multiple sources and selecting the most informative source datasets or samples through data-driven methods. For supervised statistical transfer learning methods, well-established strategies exist to select the set of informative source samples [19]. [39] introduced a prior-misalignment-risk minimizer conditional on source parameters that estimates the inclusion probability of each source dataset through a full Bayesian framework. A similar mechanism could be incorporated into the Praxis-BGM framework to reduce the risk of negative transfer.

## Supporting information

Supplementary materials are available with the manuscript submission.

## 6 Availability of data

The METABRIC breast cancer dataset [9] was accessed through `cBioPortal`[6, 10, 13], and the TCGA-BRCA dataset [26] was obtained from the R package `curatedTCGAData`[32]. The pancreas scRNA-seq benchmark data followed the article [23] and were assembled from the public GEO and ArrayExpress datasets listed in that article’s supplementary materials.

## 8 Author contributions statement

All authors contributed to the conception and development of the methodology, performed the experiments and data analyses, and jointly wrote and reviewed the manuscript.

