## Supplementary materials are available with the manuscript submission. for "Praxis-BGM: Clustering of Omics Data Using Semi-Supervised Transfer Learning for Gaussian Mixture Models via Natural-Gradient Variational Inference"

#### S.1 Superior performance of Praxis-BGM in Transferring Structural Priors on Simulated Data

To evaluate the effectiveness of leveraging informative priors by the proposed method under varying conditions, we generated data from a Gaussian mixture model with $J = 4$ components, where each component has an identity covariance matrix unless otherwise specified. Let $\mathcal{C} \subset \{1, \ldots, P\}$ denote the set of causal features and $\mathcal{N} = \{1, \ldots, P\} \setminus \mathcal{C}$ the non-causal features. For each $p \in \mathcal{C}$ and cluster $j$, $\mu_{j,p} \overset{\text{i.i.d.}}{\sim} \mathcal{N}(0, \nu)$; for $p \in \mathcal{N}$, $\mu_{j,p} = 0$ for all $j$. Unless otherwise stated, we set $\nu = 0.09$ for a low signal-to-noise (SNR) scenario, $\theta_j = 1/J$, $\mathbf{\Sigma}_j = \mathbf{I}_P \in \mathbb{R}^{P \times P}$, $P = 100$, $\mathcal{C} = 40$, and sample size $N = 100$ to generate the target data. Once the ground-truth GMM parameters for the target data are sampled, we perturbed these parameters to generate the source data with a sample size of $N_0 = 1000$, which are then used to derive the informative priors for Praxis-BGM and to train the benchmark classification-based methods.

Across four components of informative priors, we compared eight methods: (1) Linear discriminant analysis (LDA); (2) multinomial logistic regression; (3) Extreme Gradient Boosting (XGBoost), a popular machine learning classification method that leverage a large collection of weak decision trees (Chen & Guestrin 2016); (4) K-Nearest Neighbors (KNN) classifier, another machine learning classification method that predicts based on the majority class of a data point's finite nearest neighbors; and (5) Praxis-BGM with an informative prior derived from the source data. Additionally, we benchmarked these approaches with unsupervised clustering methods that do not use prior information: (6) Bayesian Gaussian Mixture Models (BGM), a VI-based GMM method using Normal-Invert-Wishart Prior without prior information (fit with $J$ fixed) ; (7) the Leiden algorithm (Traag et al. 2019), a graph-based community detection method applied to a $k$-nearest neighbor graph, which does not require $J$; and (8) Praxis-BGM with no informative prior. We applied LDA, multinomial logistic regression, KNN, and BGM implemented in *scikit-learn* with default settings (Pedregosa et al. 2011). Performance was quantified using Adjusted Rand Index (ARI), which ranges from -1 to 1 and measures concordance between estimated clusters and the simulated truth. Each scenario was replicated 500 times.

##### S.1.1 Cluster-specific Means Prior

We extensively evaluated the effect of incorporating informative priors for cluster means derived from varying source data misspecifications under the isotropic mixture of multivariate Gaussians. The mean vector to generate the source data $\boldsymbol{\mu}_j^{(0)} \in \mathbb{R}^P$ for cluster $j$ were perturbed by adding Gaussian noise $\boldsymbol{\epsilon} \sim \mathcal{N}(\mathbf{0}, 0.3^2)$ with varying proportion $\rho \in \{0\%, 10\%, 30\%, 50\%, 70\%, 100\%\}$ of features. After generating the source datasets and ground-truth cluster labels for each $\rho$ level, we computed the empirical cluster-specific means and used them as priors for Praxis-BGM to transfer knowledge to the target; we also trained supervised classification methods on the source data and applied them to predict the labels in the target data.

Figures S.1(a) show clustering/prediction performance under increasing perturbation $\rho$ applied to the source data. Praxis-BGM with mean priors achieves the highest ARI when

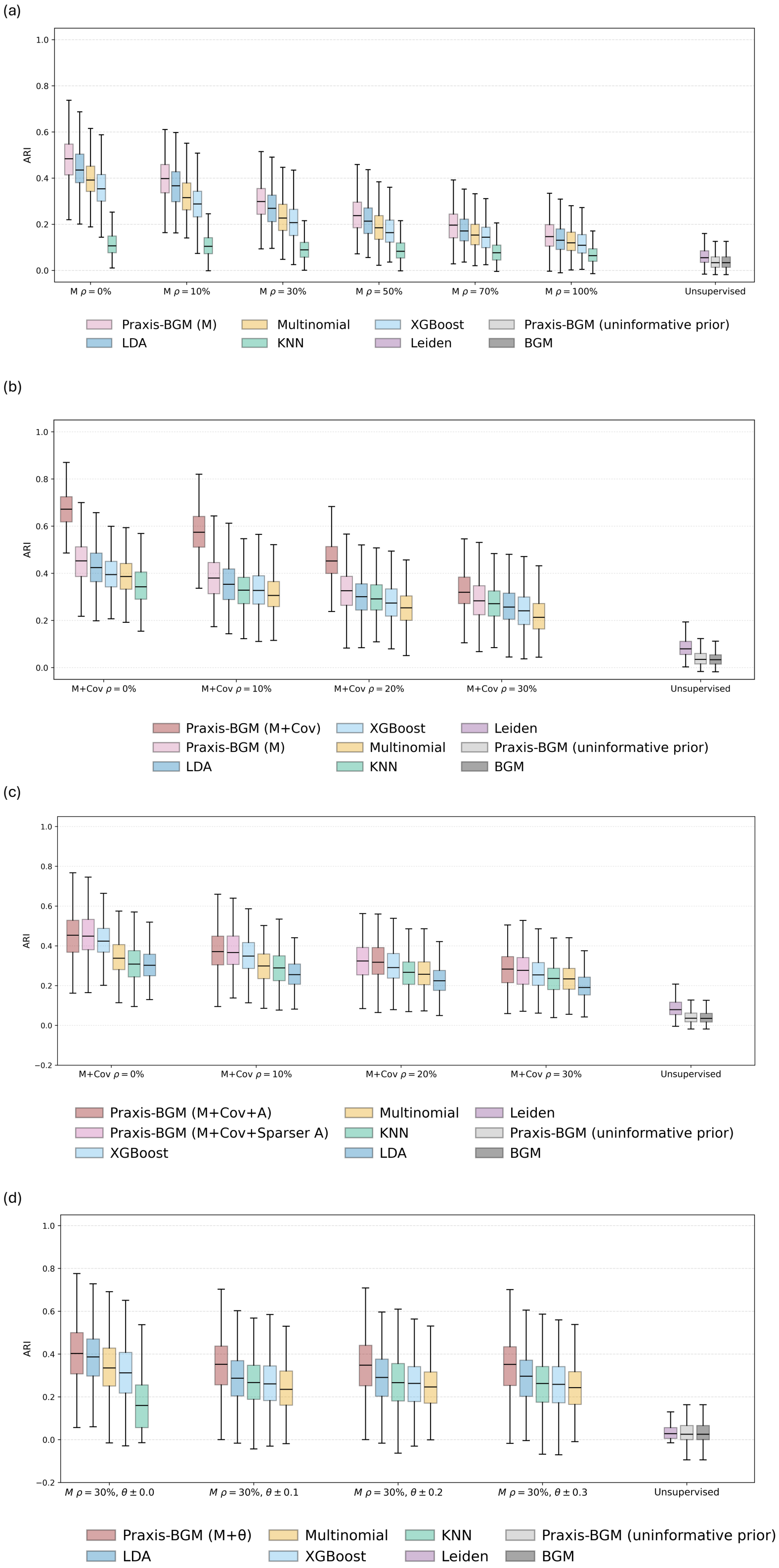


**Figure S.1:** Simulation results evaluating the effectiveness of four types of priors in Praxis-BGM under increasing levels of source-data misspecification. The y-axis shows ARI (agreement with ground-truth clusters), and the x-axis indicates misspecification severity and baseline methods. **(a)** Priors on cluster means; **(b)** Priors on means and covariances; **(c)** Priors on sparsity masks; **(d)** Priors on cluster weights.

the source data are unperturbed (i.e., when mean priors are accurate), with LDA as the next-best method. As $\rho$ increases, ARI decreases for all methods, yet Praxis-BGM continues to outperform competitors within each $\rho$ group, demonstrating robustness to cross-domain heterogeneity through its Bayesian modeling of uncertainty. All methods leveraging source information outperform unsupervised approaches (Leiden, BGM, and Praxis-BGM with an uninformative prior) even at $\rho = 100\%$, likely because Gaussian perturbations still preserve remnants of the original cluster structure. Praxis-BGM with an uninformative prior performs similarly to BGM but below Leiden, reflecting the advantage of graph-based clustering at low SNR. Overall, these results indicate that informative priors substantially improve clustering—especially when source and target data are similar or moderately distorted—and that Praxis-BGM is most successful at transferring such structural signals through mean priors.

#### S.1.2 Cluster-specific Variance-Covariance Prior

We next evaluated the impact of incorporating priors for both cluster means and variance–covariance matrices. Let $\mathbf{I}_P$ denote the $P \times P$ identity matrix and $B_j \subset \{1, \ldots, P\}$ be a contiguous block of size $\lfloor 0.1 \times P \rfloor$ chosen independently for each cluster $j$. Define the block-indicator matrix $\mathbf{E}_j \in \{0,1\}^{P \times P}$ by $(\mathbf{E}_j)_{uv} = 1$ if $u \in B_j$ and $v \in B_j$, $\quad 0$ otherwise, so that the ground-truth covariance is $\mathbf{\Sigma}_j = \mathbf{I}_P + 0.3\,\mathbf{E}_j$, yielding cluster-specific covariance patterns. As in Simulation S.1.1, we vary the perturbation level $\rho \in \{0\%, 10\%, 30\%\}$ for both $\boldsymbol{\mu}_j^{(0)}$ and $\mathbf{\Sigma}_j^{(0)}$ when generating the source data. Empirical cluster-specific means and covariances were then estimated from the source data as priors for Praxis-BGM and the source data were also used to train classification models applied to predict the target data.

Figure S.1(b) summarizes the results. With accurate priors ($\rho = 0\%$), Praxis-BGM with mean priors only (Praxis-BGM (M)) already attains a high ARI, while adding covariance priors (Praxis-BGM (M+Cov)) provides a further improvement, confirming the complementary value of variance–covariance information. As $\rho$ increases, ARI declines for all source-informed methods, yet Praxis-BGM (M+Cov) consistently performs best within each $\rho$ group. However, covariance priors are less robust to perturbation: the performance gap between Praxis-BGM (M+Cov) and Praxis-BGM (M) narrows as $\rho$ grows. Overall, when cluster separation depends on both mean and covariance structure, incorporating $\mathbf{\Sigma}^{(0)}$ enhances clustering accuracy for Praxis-BGM.

##### S.1.3 Sparsity–imposing Matrix Prior

We assessed the effect of the sparse feature–feature connectivity prior $\mathbf{A} \in \{0,1\}^{P \times P}$, in addition to mean and covariance priors, $\boldsymbol{\mu}_j^{(0)}$ and $\mathbf{\Sigma}^{(0)}$, for Praxis-BGM. $\mathbf{A}$ encodes prior independence structure, with $A_{uv} = 0$ indicating features assumed *a priori* uncorrelated. We generated $\mathbf{A}$ by selecting $\lfloor \alpha P(P-1)/2 \rfloor$ off-diagonal pairs ($\alpha = 0.05$) and defining the mask as $A_{uv} = 1$ for diagonal or selected pairs, and 0 otherwise.

As in Simulation S.1.2, we simulated four source datasets with increasing perturbation $\rho$. Target-data covariance priors were constructed as $\mathbf{\Sigma}_j^{(0)} = \mathbf{\Sigma}_j^{0*} \odot \mathbf{A}$, while the source data used unmasked covariances, so that independence information is entered only through $\mathbf{A}$. We compared Praxis-BGM using mean and covariance priors alone, with versions incorporating either the true $\mathbf{A}$ or a sparser variant $\mathbf{A}'$ formed by zeroing half of its off-diagonal entries.

Figure S.1(c) shows that Praxis-BGM performs best when accurate $\boldsymbol{\mu}^{(0)}$, $\mathbf{\Sigma}^{(0)}$, and $\mathbf{A}$ are provided, and that using $\mathbf{A}'$ yields nearly identical ARI, indicating robustness to moderate misspecification. Classification-based methods lag behind Praxis-BGM with informative priors, and unsupervised methods perform the worst. Performance remains stable across increasing $\rho$ when using $\mathbf{A}$, likely because mean and covariance priors become less influential. Overall, a well-specified sparse connectivity prior substantially enhances clustering accuracy, particularly when covariance priors contain noisy or irrelevant structure.

##### S.1.4 Cluster Weight Prior

We evaluated the effect of incorporating a cluster weight prior $\boldsymbol{\theta}^{(0)}$ under the same isotropic setting as Simulation S.1.1. The source data means were fixed at $\rho = 30\%$, and imbalanced target weights were generated via $\boldsymbol{\theta} \sim \mathrm{Dir}(0.4)$. Source weight priors were created by perturbing the true weights with uniform noise in $[-0.1, 0.1]$, $[-0.2, 0.2]$, or $[-0.3, 0.3]$ followed by a softmax transformation.

Figure S.1(d) shows that given the same mean priors ($\rho = 30\%$), Praxis-BGM with accurate weight priors achieves the highest ARI, outperforming classification-based alternatives. As weight perturbation increases, while mean-prior perturbation remains fixed at $\rho = 30\%$, ARI remains largely stable for both Praxis-BGM and classification methods. This indicates that correctly specified weight priors can provide benefits, but moderate misspecification of $\boldsymbol{\theta}^{(0)}$ does not substantially degrade performance.

#### S.2 Additional Simulation Results

##### S.2.1 Enhanced Signal-to-Noise Ratio

In the last section, we present simulation results evaluating the effectiveness of four types of priors. We also examined how Praxis-BGM withstands prior misspecification compared to benchmarking classification methods. We intentionally set the effect sizes of the causal features low by drawing them from $\mathcal{N}(0, 0.3^2)$ to create hard-to-cluster mixture data with low signal-to-noise ratio (SNR) and highlight the usefulness of the prior knowledge. In additional simulations here, we increase the effect sizes of the causal features by sampling them from $\mathcal{N}(0, 0.6^2)$ to create a higher SNR scenario, in order to assess whether Praxis-BGM continues to outperform benchmarking methods when the cluster structure in the target data is more pronounced. Figure S.2 presents the results. With a higher SNR, unsupervised methods that do not incorporate prior knowledge perform substantially better than in the low-SNR setting presented in Section S.1. Leiden achieves a higher ARI than both BGM and Praxis-BGM with uninformative priors. Interestingly, Praxis-BGM with uninformative priors outperforms BGM in simulations (b) and (c), where the target data exhibit distinct cluster structures differentiated by the covariance matrices. For other methods that leverage the source data, we observe a similar pattern to that in the low-SNR setting: Praxis-BGM makes the most effective use of prior knowledge and consistently outperforms classification-based methods across all types of priors, including cluster-specific means, variance–covariance matrices, the sparsity-imposing matrix, and cluster weights. Additionally, perturbations to the source data appear to have less impact on clustering performance; as $\rho$ increases, we do not observe a significant drop in median ARI for Praxis-BGM, although the variance of ARI becomes slightly larger.

##### S.2.2 Batch Effects

It is also of interest to examine noise or heterogeneity in the source data relative to the target that is not random at the cluster level but instead represents a consistent shift across all clusters, mimicking a batch effect between two datasets. This is a common scenario in omics studies, where different laboratories or experimental batches often introduce baseline differences. Figure S.3 presents the simulation results for mean and variance–covariance priors when the source data are perturbed only by a batch effect, under a low SNR setting. Across Figure S.3(a) and (b), we observe that within each source data group, Praxis-BGM outperforms other classification-based methods due to the nature of Bayesian inference and its ability to quantify uncertainty. This is consistent with the previous findings. However, as $\rho$ increases in Figure S.3(a), the decreasing trend in ARI is less pronounced in comparison to

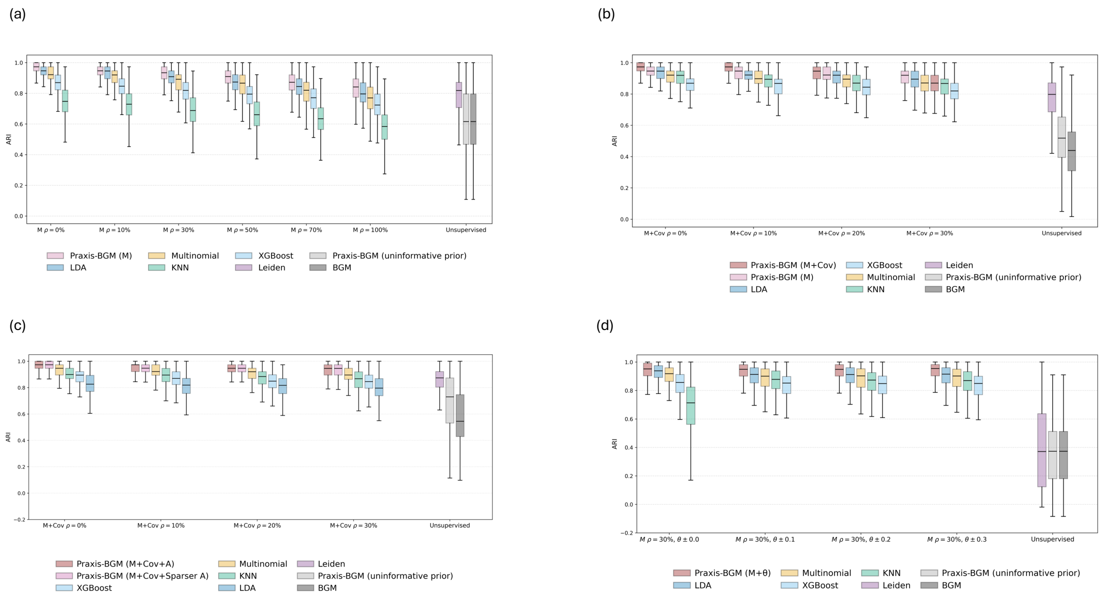


Figure S.2: Simulation results evaluating the effectiveness of the four components of the priors in Praxis-BGM under increasing levels of source-data misspecification in the high-SNR setting. The y-axis shows ARI (agreement with ground-truth clusters), and the x-axis represents increasing levels of misspecification along with unsupervised baseline methods. **(a)** Informative priors on cluster means; **(b)** Informative individual and joint priors on cluster means and structured variance–covariance matrices; **(c)** Priors on the sparsity-imposing matrix; **(d)** Priors on cluster weights.

the scenarios presented in Section S.1. In Figure S.3(b), the decrease is even less pronounced when mixture data are also driven by covariance patterns. These results suggest that Praxis-BGM can effectively accommodate batch effects as it incorporates information from the target data. Systematic shifts preserve relative cluster separations and enable the model to leverage structural information encoded in the mean and variance-covariance priors despite global biases. Overall, these findings indicate that informative mean and variance-covariance priors substantially improve clustering performance when approximately correct, and that Praxis-BGM remains robust to batch-effect-like distortions as long as the underlying relational structure is preserved.

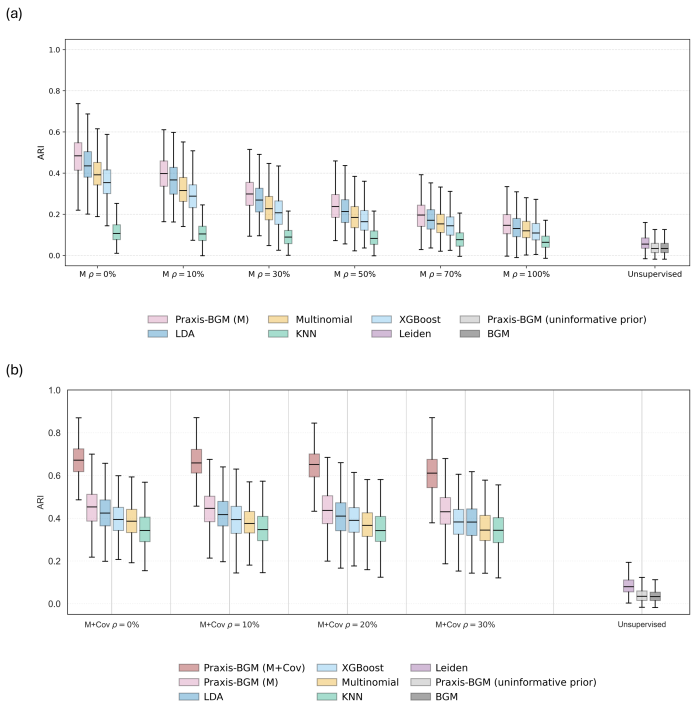


Figure S.3: Simulation results evaluating the effectiveness of mean and variance–covariance priors for Praxis-BGM on clustering performance when the source data are perturbed only by a batch effect, under a low-SNR setting. **(a)** Informative priors on cluster means. **(b)** Informative priors on both cluster means and structured variance–covariance matrices.

##### S.2.3 Computation Time with Different Hardwares and Scalability

To assess computational scalability, we use the simulation setup from Section S.1 but with dimensionality ($P \in \{50, 100, 500\}$), sample size up to $N = 50{,}000$, and cluster numbers ($J \in \{4, 16\}$). These settings emulate large-scale omics data applications, where both algorithmic efficiency and stability are critical. In theory, the total time complexities of both BGM and Praxis-BGM are $\mathcal{O}(NJP^2)$ because of the cost of updating and inverting cluster-specific covariance matrices, while the time complexity of Leiden is $\mathcal{O}(NJ)$, as it is graph-based. Since Praxis-BGM is implemented in JAX (Bradbury et al. 2018), which provides a NumPy-style interface capable of GPU acceleration, we ran Praxis-BGM on a T4 GPU, while the baseline methods (BGM and Leiden) were run on CPUs. Here, we focus on the computation time required for convergence for each method, as clustering performance has already been evaluated in Section S.1.1. We replicated each analysis 50 times.

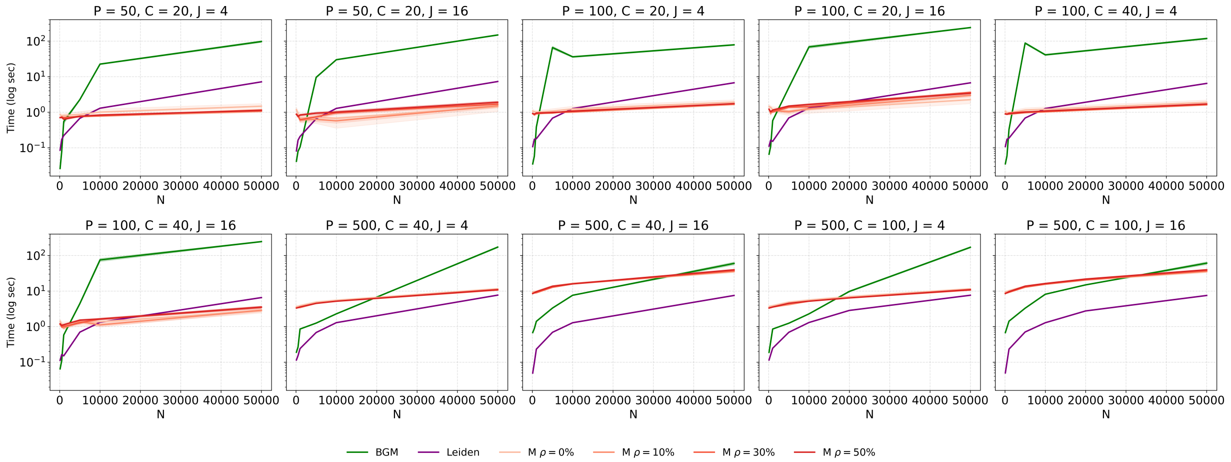


Figure S.4: Computation time (seconds, log scale) across 10 scenarios. Run on a T4 GPU, Praxis-BGM demonstrates linear scalability with $N$ and robustness to increasing feature dimension $P$ and number of clusters $J$.

Figure S.4 shows the median computation time (reported to reduce the influence of potential outliers for baseline methods) on a log scale. Different prior specifications of Praxis-BGM result in similar computation times, although in some cases, more accurate priors can lead to faster convergence. Praxis-BGM scales approximately linearly with $N$ and remains efficient as $P$ and $J$ increase, with log runtime between 1 and 10 even for $N = 50{,}000$ and $P = 500$. For lower dimensions ($P = 50, 100$), the log runtime is about 1 regardless of $N$ and $J$, reflecting the initialization cost of the model, after which inference proceeds with negligible additional computation. In contrast, BGM's EM-based inference exhibits steeper growth in runtime with both $N$ and $J$ at all dimensionalities. As expected, Leiden scales more favorably than BGM and Praxis-BGM in $N$ at higher dimensions (e.g., $P \geq 100$), but for $P = 50$ and $P = 100$, its runtime increases with $N$ and exceeds that of Praxis-BGM. Therefore, when the feature dimension is manageable (e.g., $P \leq 100$), Praxis-BGM is more computationally efficient than Leiden and remains as fast as the sample size $N$ increases. This makes Praxis-BGM well-suited and scalable for clustering large-scale scRNA-seq datasets or dimension-reduced/feature-selected omics data. These results demonstrate that Praxis-BGM's natural-gradient variational inference and the use of the JAX framework for just-in-time (JIT) compilation provide stable estimation and scalability for large datasets with approximate linear time cost, even though the complexity of still $\mathcal{O}(NJP^2)$.

We then investigated the computation time of Praxis-BGM on the same large-scale data when run on different hardware platforms (CPU, GPU, and TPU), since the JAX framework provides optimized acceleration across all of them. We set $N = 10{,}000$, feature dimension $P = 100$, and number of clusters $J = 4$. After 50 replications, we computed the mean computation time for each Prxis-BGM with different prior specifications on all hardware. Figure S.5 presents the results. We observe minimal differences across prior specifications within each hardware type, but a large discrepancy between CPU and GPU/TPU runtimes for the same prior. Running Praxis-BGM on a V5e-1 TPU yields the fastest computation

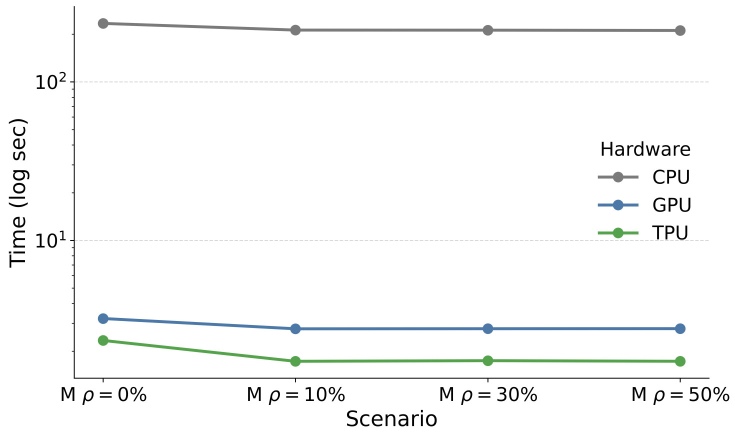


Figure S.5: Computation time (seconds, log scale) for Praxis-BGM across CPU, GPU, and TPU hardware under different prior qualities. Results are based on simulated data with $N = 10{,}000$, feature dimension $P = 100$, and number of clusters $J = 4$.

across all priors, with the T4 GPU slightly slower but comparable, while CPU execution is substantially slower. Thus, for large-scale applications of Prxis-BGM, such as label transfer for scRNA-seq Data, we recommend using a TPU or a GPU to improve efficiency.

### S.3 Additional Analyses for the Breast Cancer Bulk Transcriptomics Application

#### S.3.1 Clustering-Driven Gene Selection via Bayes Factors

To assess whether feature $p$ contributes to clustering, we compare its marginal likelihood under the posterior GMM to that under a homogeneous mixture null. Let $\mathcal{D}_p = \{D_{ip}\}_{i=1}^N$ denote the observed values of feature $p$. Under the null, all components share the same parameters, i.e., $\mu_{jp} = \hat{\mu}_p$ and $\sigma^2_{jp} = \hat{\sigma}^2_p$ for all $j$, with equal weights $1/J$, so that no cluster-specific structure is present (equivalently, the null model collapses to a single Gaussian).

We define an approximate Bayes factor (BF) as a plug-in ratio of marginal likelihoods, under the assumption of equal prior probabilities:

$$\mathrm{BF}_p = \exp\left\{\sum_{i=1}^{N} \log\left[\frac{\sum_{j=1}^{J} \theta_j \, \mathcal{N}(D_{ip} \mid \mu_{jp}, \sigma_{jp}^2)}{\mathcal{N}(D_{ip} \mid \hat{\mu}_p, \hat{\sigma}_p^2)}\right]\right\}. \tag{S.3.1}$$

This quantity provides an empirical approximation to the BF, and $\mathrm{BF}_p$ can also be interpreted as the ratio of the posterior probabilities for feature inclusion. Interpretation of $\mathrm{BF}_p$ follows the conventional evidence categories of Jeffreys' scale (Jeffreys 1961). To identify genes driving breast cancer subtype differentiation, we computed BFs using the above method for each gene using the posterior Praxis-BGM model fitted to TCGA-BRCA RNA-seq data with METABRIC-derived priors.

#### S.3.2 Sensitivity Analysis of Clustering and Classification Performance across Different Numbers of Top HVGs

For the TCGA-BRCA bulk transcriptomics transfer-learning application from METABRIC, we performed an additional sensitivity analysis to evaluate how the number of selected highly variable genes (HVGs) influences clustering and classification performance.

Although PAM50 genes are directly related to the ground-truth subtype labels used for evaluation, we did not consider a PAM50-restricted analysis to reflect the intended use case. In this illustrative application, PAM50 subtype serves primarily as an external reference for validation, whereas the goal of Praxis-BGM is to recover meaningful structure from a broader and less curated transcriptomic feature space. This setting is more realistic for applications in which validated marker or causal gene sets are unknown.

To identify HVGs, we firstly subsetted METABRIC and TCGA-BRCA to the shared gene set ($p$ = 16,151) and ranked genes by expression variance in the source dataset, METABRIC, after $\log_2$ transformation. HVGs were selected using the source data alone, rather than after concatenating source and target datasets, because domain heterogeneity can distort feature variability and we assume that the source cohort is sufficiently large to provide a stable representation of gene-level variation in a Bayesian transfer learning setting.

We examined the ranked variance spectrum of the top 5000 shared genes in METABRIC (Figure S.6), which roughly showed an elbow around the top 1000 genes. In the main analysis, we used the top $p = 1000$ HVGs, relying on the elbow method on the ranked gene variance plot to remove genes with low-information. Including substantially more than 1,000 genes may contribute limited additional signal while increasing noise and computational burden, whereas including fewer genes may result in loss of information; therefore, a tradeoff is involved. To examine it, we therefore evaluated $p = 500$, 1000, and 2000 HVGs in a sensitivity analysis. We benchmark Praxis-BGM with informative priors on the source data with BGM, LDA, multinomial logistic regression (MultinomialReg), and XGBoost (XGB).

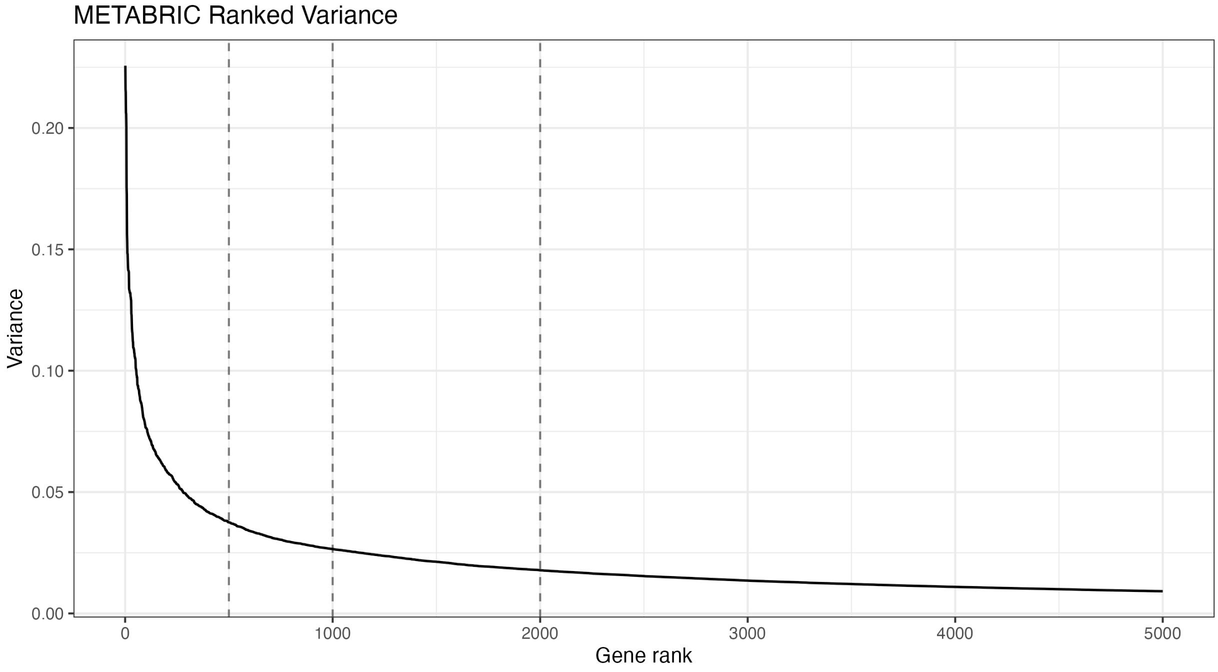


Figure S.6: Ranked gene expression variance in METABRIC (shared genes across domains).

As shown in Figure S.7, performance as measured by ARI on the 500 TCGA-BRCA subjects with available subtype labels was generally stable across feature settings for most methods.

Interestingly, LDA was a notable exception, showing a pronounced decrease at 2000 genes, which suggests sensitivity to additional noisy features for this method. Praxis-BGM achieved the highest ARI across all three settings, with its best performance at 500 HVGs (ARI = 0.471) and only modest decreases at 1000 HVGs (ARI = 0.430) and 2000 HVGs (ARI = 0.426).

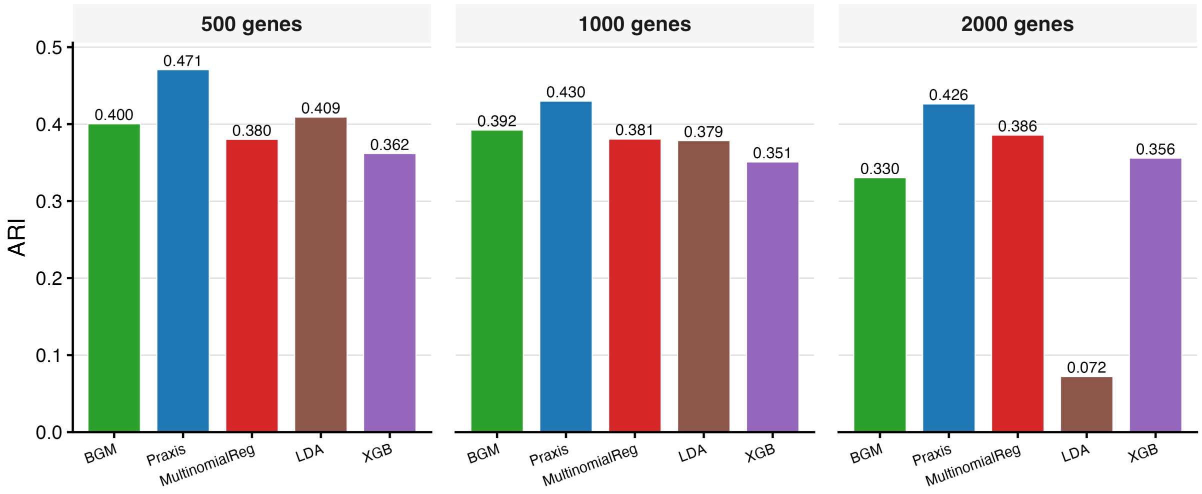


Figure S.7: ARI with PAM50 ground-truth subtype comparison across benchmark methods using 500, 1000, and 2000 HVGs.

Overall, these results suggest that including genes beyond the variance elbow may reduce performance for some methods such as LDA, whereas Praxis-BGM remains relatively robust. Although Praxis-BGM attained its highest ARI at 500 HVGs, we used $p = 1000$ HVGs in the main analysis as we would like to stay with the variance-based filtering approach to select the number of top HVGs instead of the downstream performance-based analysis. In practice, ground-truth subtype labels for validation may not be available, so the number of HVGs often must be determined solely from the variance profile. In addition, we were also interested in downstream survival analysis, for which restricting the feature set to 500 genes may exclude potentially informative variation.

##### S.3.3 Performance Evaluation in Extreme HDLSS Settings

To better assess the performance of Praxis-BGM in a extreme high dimensionality and limited sample sizes (HDLSS) setting, we conducted a sensitivity analysis for the TCGA-BRCA bulk transcriptomics application by repeatedly subsampling the target cohort to $n = 25$ and $n = 50$. Although TCGA-BRCA contains 1,060 samples in total, agreement between estimated clusters and ground-truth subtype labels could only be evaluated on the subset of 500 subjects with available subtype annotations. We therefore based this analysis on repeated subsamples from this labeled subset.

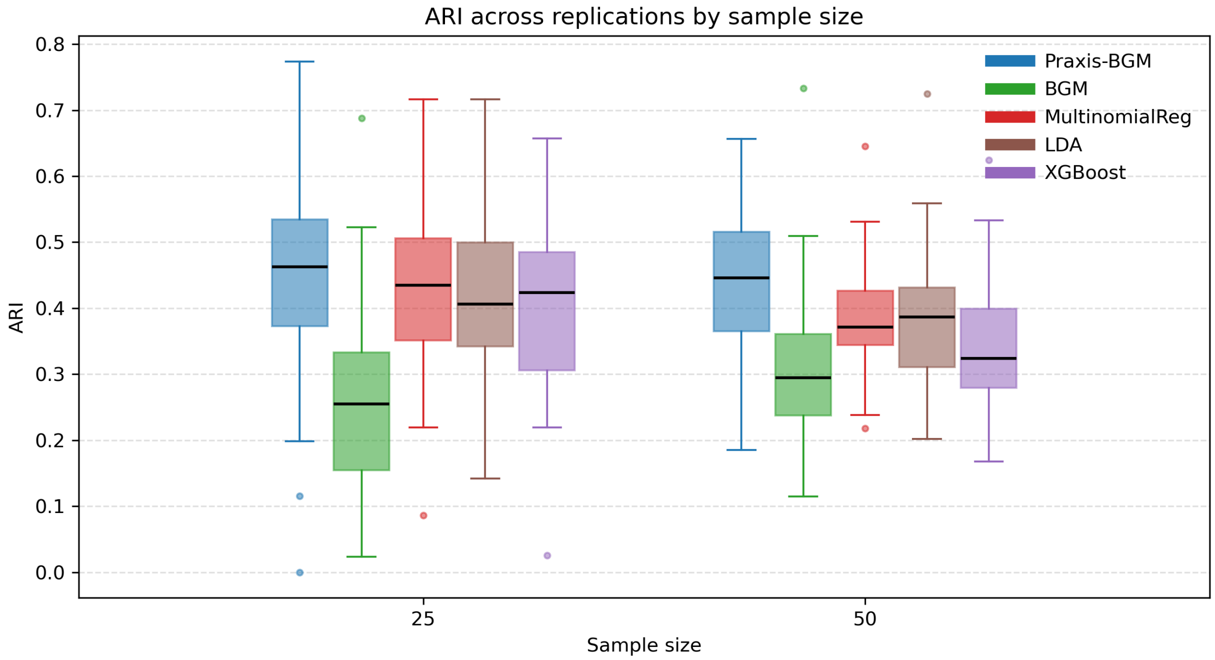


Figure S.8: ARI with PAM50 ground-truth subtype comparison across methods under small target sample sizes in TCGA-BRCA with 30 replications

Figure S.8 summarizes ARI with PAM50 ground-truth subtype across 30 replications for Praxis-BGM and competing clustering and classification methods. Across both sample sizes, Praxis-BGM achieved the highest median ARI relative to GMM, BGM, multinomial logistic regression, and XGBoost. These results support the intended use of Praxis-BGM in more challenging HDLSS regimes, where target sample sizes are very limited and the dimensionality is relatively high ($p = 1000$).

The advantage of Praxis-BGM arises not only from incorporating informative priors derived from the source dataset, but also from updating the target-domain model through variational inference, allowing the mixture model to adapt to target-specific structure even when only a small number of target samples are available. This combination appears to offer an advantage over directly applying deterministic classification models to the target data, and also over purely unsupervised methods, for which meaningful cluster structure can be difficult to estimate reliably in very small samples. One limitation of this analysis is that it is restricted to the 500 labeled TCGA-BRCA subjects and therefore does not fully reflect performance on the complete cohort of 1,060 samples. In particular, the remaining unlabeled subjects could potentially provide additional information for posterior updating in Praxis-BGM.

##### S.3.4 Data Preprocessing for In-Serial Bayesian Clustering on the miRNA Layer of TCGA-BRCA Using Praxis-BGM

After estimating Praxis-BGM posterior clusters ($N = 1060$) on the transcriptomic (bulk RNA-seq data) layer of TCGA-BRCA, which were informed by priors from the source data METABRIC to promote transfer learning, we used the posterior RNA Praxis-BGM clusters to construct cluster-specific priors for estimating the miRNA layer of the same subjects. This enabled a second, in-serial Bayesian update on the miRNA data in TCGA-BRCA. As miRNA measurements were available for only 731 of the 1,060 subjects, the posterior RNA Praxis-BGM cluster labels were correspondingly subset to these individuals for this subsequent analysis. After quality control to remove missing miRNA features or features with all zero-count entries, there were 1,002 mature miRNA features remaining. We next applied $\log_2$ transformation to the miRNA data and used the elbow of the ranked variance profile as a data-driven heuristic to exclude lower-variance, potentially less informative miRNA features. We then standardized the selected top 100 most variable miRNA features. Conditioned on the RNA Praxis-BGM cluster with subtype annotations, we computed empirical means and covariance matrices in the miRNA feature space to construct informative priors. The resulting priors were then supplied to Praxis-BGM to infer posterior miRNA clusters, which were subsequently evaluated by their ARI with PAM50 labels among subjects with available annotations and by comparison with the RNA-based Praxis-BGM clusters to assess whether incorporating miRNA information provides additional benefit.

#### S.4 Additional Analyses for the Cross-Platform Label Transfer Application in Pancreatic scRNA-seq

##### S.4.1 Sensitivity Analysis of DEG-Based Feature Selection for Cross-Platform Pancreatic scRNA-seq Label Transfer Benchmark Methods

As an additional sensitivity analysis for the label transfer analysis using Praxis-BGM and benchmark methods across pancreatic scRNA-seq platforms presented in the main manuscript, we investigated the performance of KNN-based benchmark label transfer methods from using a smaller and more targeted set of source-derived marker genes or differentially expressed genes (DEGs), rather than the original workflow based on 2,000 highly variable genes (HVGs). The goal is to examine the hypothesis that KNN-based methods may perform more favorably when trained on biologically informative features instead of a broader transcriptomic feature space.

We considered two DEG-based feature selection strategies using the reference dataset. First, we identified up to the top 100 significant DEGs for each reference cell type and used the union of these genes, resulting in 917 features. Under this setting, the performance of all methods was essentially unchanged relative to the original 2,000-HVG workflow in the main manuscript, suggesting that a moderately reduced but still broad marker set preserves most of the relevant signal.

**Table S.1:** Benchmarking label transfer methods on pancreatic single-cell transcriptomic data using the top-10 DEGs-per-cell-type union feature selection strategy (130 genes), evaluated against expert annotations.

| **Method** | **ARI** | **Accuracy** | **Embedding** |
|---|---|---|---|
| Praxis-BGM | 0.943 | 0.965 | PCs |
| ComBat-seq + KNN | 0.932 | 0.957 | PCs |
| Scanorama + KNN | 0.929 | 0.945 | PCs |
| Scanpy-ingest | 0.905 | 0.922 | PCs |
| scANVI | 0.886 | 0.906 | scVI |

We then considered a more aggressive feature restriction by selecting up to the top 10 significant DEGs for each cell type, yielding a total of 130 genes for label transfer. Under this more targeted feature set, we presented in the results in Table S.1. Scanorama + KNN and `Scanpy-ingest` showed improved performance relative to the original HVG-based analysis, with increases in accuracy (Scanorama + KNN increases from 0.916 to 0.945; `Scanpy-ingest` increases from 0.915 to 0.922). ComBat-seq + KNN remained highly competitive and changed only marginally. Praxis-BGM showed a modest reduction in performance (ARI: 0.957 to 0.943; accuracy: 0.975 to 0.965), but notably still achieved the highest ARI and highest accuracy among all compared methods under this restricted-feature scenario. In contrast, scANVI exhibited a larger decrease (ARI: 0.912 to 0.886; accuracy: 0.924 to 0.906).

These findings highlight an important distinction between method classes. ComBat-seq + KNN, Scanorama + KNN, and `Scanpy-ingest` intrinsically operate as deterministic KNN-based classifiers and may therefore benefit from a sparse, highly discriminative marker set. By contrast, Praxis-BGM relies on clustering, and scANVI relies on integrated latent-space learning, where broader HVG-based inputs may better preserve global transcriptomic structure and relationships among closely related cell types. Importantly, even under aggressive DEG restriction, the prior-regularized clustering strategy of Praxis-BGM remained the strongest-performing approach overall.

##### S.4.2 Label Transfer Across Platforms in Pancreatic scRNA-seq Data with Fewer Reference Cell Types

In the label transfer analysis using Praxis-BGM across pancreatic scRNA-seq platforms presented in the main manuscript, the annotated reference subset included T cells, whereas the query subset did not. In this analysis, we used the same reference and query datasets as in the main manuscript but removed cells annotated with rare cell types, including "Schwann (0.152%)," "Epsilon (0.210%)," "T cell (0.0817%)," and "Mast (0.292%)" to emulate a scenario in which the reference contained fewer cell types than the query and create a label transfer task with a misspecified reference dateset (Figure S.9(a)). Note that the T cell does not exist in the ground-truth cell type annotation in the query dataset. We compared Praxis-BGM with benchmarking methods, and the results are presented in Table S.2.

Overall, Praxis-BGM achieved the highest ARI and accuracy (97.1% accuracy, ARI = 0.955), nearly matching its superior performance in the main manuscript, where the reference dataset was fully comprehensive. ComBat-seq + KNN also substantially outperformed other methods except Praxis-BGM in terms of ARI and accuracy (95.1% accuracy, ARI = 0.932),

**Table S.2:** Benchmarking label transfer methods on Pancreas single-cell transcriptomic data against expert annotations.

| Method | Time (s) | ARI | Accuracy | Hardware | Embedding |
|---|---|---|---|---|---|
| Praxis-BGM | 2.95 | **0.955** | **0.971** | T4 GPU | PCs |
| ComBat-seq + KNN | 4.46 | 0.932 | 0.951 | CPU | PCs |
| Scanorama + KNN | 9.61 | 0.910 | 0.912 | CPU | PCs |
| Scanpy-ingest | 7.31 | 0.898 | 0.912 | CPU | PCs |
| scANVI | 30.0 | 0.914 | 0.923 | T4 GPU | scVI |

maintaining a similar performance as in the main manuscript setting, though still falling short of Praxis-BGM. Overall, all methods degraded slightly due to the deletion of rare cell types in the reference data compared to the performance in the main manuscript. In terms of computation time, all methods showed runtimes consistent with their performance in the main manuscript. Figure S.9(b) shows the confusion heatmap comparing predicted cell types to the ground-truth annotations in the query data. The apparent misalignment for the last three cell types is expected, as these cell types are absent from the reference and therefore cannot be predicted by any of the methods. For the remaining cell types, the pattern across methods is consistent with what we observed when using the complete reference data.

These results further highlight the robustness of Praxis-BGM for label transfer across datasets generated from different technology platforms in a more realistic setting (the reference dataset is not comprehensive enough). Whether or not the reference dataset contains a comprehensive set of cell types, Praxis-BGM effectively leverages the available prior information to estimate the most accurate assignments through variational inference.

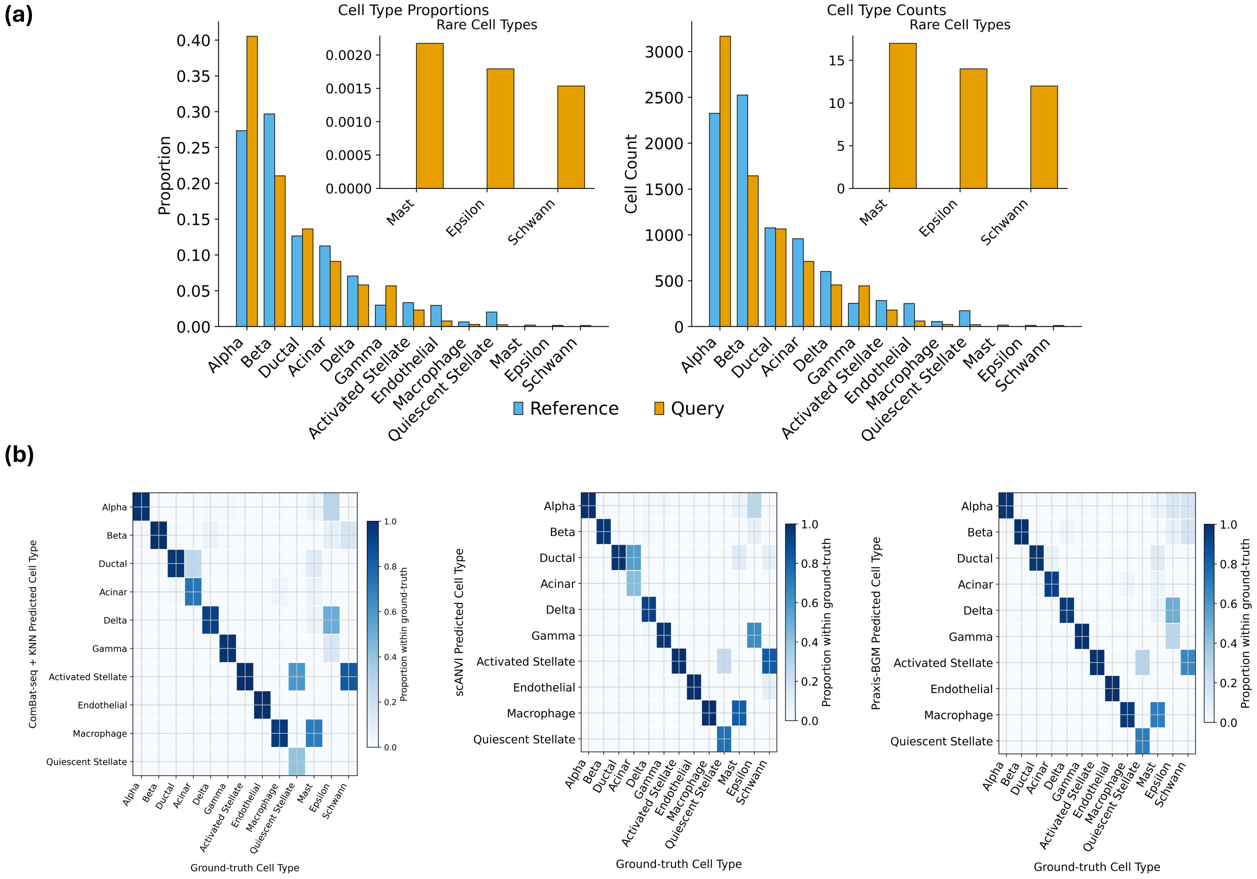


**Figure S.9:** (a) Ground-truth cell type proportions (left) and count (right) distributions for the reference and query datasets. (b) Confusion heatmap comparing predicted cell types to ground-truth annotations in the query dataset for ComBat-seq + KNN, scANVI, and Praxis-BGM, with color intensity indicating the within–cell type proportion.
